# Thalamo-Prefrontal Connectivity Correlates with Early Command Following After Severe Traumatic Brain Injury

**DOI:** 10.1101/2022.01.06.475131

**Authors:** Megan E. Cosgrove, Jordan R. Saadon, Charles B. Mikell, Patricia L. Stefancin, Leor Alkadaa, Zhe Wang, Sabir Saluja, John Servider, Bayan Razzaq, Chuan Huang, Sima Mofakham

## Abstract

Recovery of consciousness after traumatic brain injury (TBI) is heterogeneous and difficult to predict. Structures such as the thalamus and prefrontal cortex are thought to be important in facilitating consciousness. We sought to investigate whether the integrity of thalamo-prefrontal circuits, assessed via diffusion tensor imaging (DTI), was associated with the return of goal-directed behavior after severe TBI. We classified a cohort of severe TBI patients (N = 25, 20 males) into *Early* and *Late/Never* outcome groups based on their ability to follow commands within 30 days post-injury. We assessed connectivity between whole thalamus, and mediodorsal thalamus (MD), to prefrontal cortex (PFC) subregions including dorsolateral PFC (dlPFC), medial PFC (mPFC), anterior cingulate (ACC), and orbitofrontal (OFC) cortices. We found that the integrity of thalamic projections to PFC subregions (L OFC, L and R ACC, and R mPFC) was significantly associated with *Early* command-following. This association persisted when the analysis was restricted to prefrontal-mediodorsal (MD) thalamus connectivity. In contrast, dlPFC connectivity to thalamus was not significantly associated with command-following. Using the integrity of thalamo-prefrontal connections, we created a linear regression model that demonstrated 72% accuracy in predicting command-following after a leave-one-out analysis. Together, these data support a role for thalamo-prefrontal connectivity in the return of goal-directed behavior following TBI.

## Introduction

Patients with severe traumatic brain injury (TBI), defined as initial Glasgow Coma Scale (GCS) < 8 and loss of consciousness for at least 24 hours, recover consciousness at variable intervals (Winans et al. 2019; Wang et al. 2021). The mechanisms underlying coma and recovery of consciousness in patients with TBI are not fully understood. The combined effects of both primary injury to brainstem arousal nuclei (Edlow et al. 2013; Valko et al. 2016) and corticothalamic circuits (Forgacs et al. 2017), and secondary injury due to ischemia (Stein et al. 2004), microvascular dysfunction (Glushakova, Johnson, and Hayes 2014), and edema (Marmarou et al. 2006), affect the level of consciousness and ultimately patient outcomes after TBI.

After the acute phase of injury has passed, approximately 70% of patients eventually regain consciousness (Nakase-Richardson et al. 2012). Clinical assessment of consciousness generally involves testing command-following ability and other indicators of brain function (Teasdale and Jennett 1974; Giacino, Kalmar, and Whyte 2004; Wijdicks et al. 2005). While consciousness may precede the ability to follow commands, recovery of command-following is an important predictor of functional outcome (Suskauer et al. 2009; Greenwald et al. 2015). Importantly, the time until recovery of command-following is also a robust predictor of outcome (Whyte et al. 2001; Zhao et al. 2021). However, the underlying mechanisms that enable command-following are not well understood.

Although consciousness is required for voluntary behavior, less is known about what brain circuits support the language-guided control needed to follow verbal commands. The absence of this voluntary behavior, however, is not a prerequisite for consciousness, as a subset of these patients may display cognitive-motor dissociation (Owen et al. 2006). Resting fMRI connectivity of frontal networks differentiated brain hemorrhage patients who followed commands from those who did not (Mikell et al. 2015). Invasive recordings of the prefrontal cortex also revealed the emergence of complex cortical activity as patients recovered consciousness and eventually followed commands (Mofakham et al. 2021). Small randomized trials support a role for stimulation of prefrontal cortex to improve command-following behavior; a third of patients who responded to transcranial direct current stimulation (tDCS) of PFC regained command-following ability (Thibaut et al. 2014). The patients who responded to tDCS tended to have preserved PFC and thalamic gray matter (Thibaut et al. 2015).

The role of the frontal lobes in facilitating consciousness remains controversial (Boly et al. 2017; Odegaard, Knight, and Lau 2017). The prefrontal cortex is thought to be part of a large-scale “ignition” network that is activated when stimuli become conscious (S. Dehaene et al. 2001; Melloni et al. 2007; Stanislas Dehaene and Changeux 2011). However, data from newer “no-report” paradigms has challenged this view (Frässle et al. 2014), and investigators have suggested that prefrontal activation simply reflects the behavioral report of conscious awareness. Given the prefrontal cortex’s role in encoding rule-guided behavior (J. D. Wallis, Anderson, and Miller 2001; Jonathan D. Wallis and Miller 2003), action monitoring (Gehring and Knight 2000; Walton, Devlin, and Rushworth 2004), and cognitive control (Shenhav, Cohen, and Botvinick 2016), it seems likely that the prefrontal cortex is essential to the recovery of language-guided behavior after TBI.

How the thalamus facilitates PFC function is a subject of active investigation (Sébastien Parnaudeau, Bolkan, and Kellendonk 2018). Traditionally, the thalamus was known as a passive relay of sensory signals to the cortex. However, recent studies suggest that the thalamus, and thalamocortical connectivity specifically, are also critically important for consciousness (Monti et al. 2015; Zheng et al. 2017). Severe TBI results in injuries to grey matter in the cortex and thalamus as well as widespread damage to subcortical white matter (Anderson et al. 1996; Newcombe et al. 2007; Moen et al. 2014). In a cohort of these patients, thalamic atrophy was predictive of prolonged unconsciousness (Lutkenhoff et al. 2013). Furthermore, structural and functional connectivity between thalamic and prefrontal regions has been shown to correlate with the level of consciousness (Fernández-Espejo et al. 2011; Monti et al. 2015). Thalamic stimulation has also demonstrated promise in augmenting the level of consciousness in both animal models (Redinbaugh et al. 2020; Bastos et al. 2021) and TBI patients (Schiff et al. 2007; Cain et al. 2021).

Higher-order thalamic nuclei such as the mediodorsal nucleus (MD) are actively involved in sustaining and switching task-related cortical representations by gating cortico-cortical connectivity (Schmitt et al. 2017). MD is the largest thalamic output to PFC and is critical for several cognitive tasks such as working memory (Peräkylä et al. 2017). Several studies have revealed that MD activity is the key to sustaining PFC information during the delay period of working memory tasks (Alexander and Fuster 1973; Schmitt et al. 2017). Moreover, MD lesions lead to deficits in cognitive control (Dolleman-van der Weel, Morris, and Witter 2009; Sebastien Parnaudeau et al. 2013). While MD’s exact function is not yet clear, recent data support the view that MD signals the representing behavioral context and dynamically controls the gain of cortico-cortical connections to allow for the formation of transient neuronal ensembles (Schmitt et al. 2017) and suppress competing motor plans (Rikhye, Gilra, and Halassa 2018) in response to task demands.

We, therefore, hypothesized that thalamic input (in particular MD) to PFC facilitates the dynamic formation of neuronal assemblies required for recovery of consciousness and command-following after severe TBI. Diffusion MRI (dMRI) is a modality that can characterize abnormalities to these thalamo-prefrontal projections by detecting the restricted diffusion of water in the brain. By reconstructing the white-matter tracts of interest, quantitative measurements can shed light on the differences in tissue microstructure that may prognostically predict the loss or recovery of consciousness among TBI patients (Bigler 2010). To test our hypothesis, we assessed the integrity of thalamocortical projections on dMRI of TBI patients and developed a model for predicting command-following based on tractography analysis of these connections.

## Methods

### Ethics Statement

This retrospective study was approved by the Stony Brook University Hospital (SBUH) Committee on Research Involving Human Subjects (CORIHS) with a waiver of consent (2019-00464).

### Data Availability

All scans and clinical details (with identifying information removed) are available to interested investigators upon reasonable request to the corresponding author.

### Study Subjects

Adult TBI patients (age ≥ 18) were screened for severe injury by searching chart documentation of initial Glasgow Coma Scale (GCS) < 8 or decompensation to GCS < 8 during the initial admission. Data and imaging were collected retrospectively from all patients who met the study’s criteria (N = 25). Clinical information collected from patients included the date of injury, date of initiation of command-following, initial GCS score, findings on imaging, and interventions performed (**Table 1**). We identified the first day on which patients followed a simple verbal command (i.e., toe wiggling or tongue protrusion). Patients were subsequently categorized into the *Early group* (command-following within 30 days) and *Late/Never group* (after 30 days or never/death).

**Table 1.** Clinical Characteristics of patients.

| Subject Number | Age | Gender | Initial GCS | Days to Command Follow | Mechanism of Injury | Imaging Findings | Intervention |
| --- | --- | --- | --- | --- | --- | --- | --- |
| 1 | 25 | M | 8 | 6 | Motorcycle | SAH | EVD |
| 2 | 57 | M | 7 | 16 | Fall | SAH, SDH | EVD, ICP |
| 3 | 35 | M | 3 | 58 | Fall | SDH, occipital skull fracture | Craniotomy, EVD, ICP |
| 4 | 70 | M | 7 | 10 | Fall | Contusion, IVH |  |
| 5 | 84 | M | 4 | 3 | Found Down | SAH, contusion | Burr hole craniostomy |
| 6 | 24 | M | 7 | 12 | Motorcycle | SDH, IVH, contusion | Craniotomy, ICP |
| 7 | 25 | F | 3 | 11 | MVA | SDH, contusion | Craniotomy |
| 8 | 61 | M | 6 | 2 | MVA | IVH, contusion |  |
| 9 | 21 | F | 7 | 21 | MVA | SAH, SDH, contusion | ICP |
| 10 | 84 | M | 3 | 53 | Fall | IVH, contusion | EVD |
| 11 | 77 | M | 7 | 6 | Fall | SDH | Craniotomy |
| 12 | 61 | M | 3 | 24 | Bicycle | SAH, contusion, frontal skull and orbital fractures | EVD, ICP |
| 13 | 76 | M | 7 | Never | Fall | SDH | Craniotomy |
| 14 | 31 | M | 8 | 3 | Ped struck | Soft tissue swelling | ICP |
| 15 | 30 | M | 13 | 3 | Assault | SDH, SAH, contusion |  |
| 16 | 63 | M | 7 | 2 | Fall | SDH | Craniotomy |
| 17 | 48 | M | 3 | 35 | Ped struck | SAH, SDH | Craniotomy, EVD, ICP |
| 18 | 44 | F | 5 | 6 | MVA | SAH, orbital roof fracture | ICP |
| 19 | 30 | M | 3 | 13 | Motorcycle | Intraparenchymal hemorrhage | ICP |
| 20 | 43 | M | 3 | 178 | Bicycle | SAH | ICP |
| 21 | 68 | M | 3 | 15 | Ped struck | SAH, SDH |  |
| 22 | 27 | M | 3 | 144 | Ped struck | SAH, SDH | Craniotomy, EVD |
| 23 | 74 | F | 8 | Never | MVA | SAH, SDH | Craniotomy, ICP |
| 24 | 34 | F | 7 | 127 | MVA | SAH, SDH, facial bone fractures | ICP |
| 25 | 57 | M | 3 | Never | Bicycle | SAH, SDH, contusion | ICP |

SAH, subarachnoid hemorrhage; SDH, subdural hematoma; IVH, intraventricular hemorrhage; EVD, extraventricular drain; ICP, intracranial pressure monitor

### MRI and DWI acquisition

Clinical teams obtained MRIs for clinical prognosis 2–50 days post-injury, with a mean period of 11 ± 9 days. All images were acquired on the same 3 Tesla Siemens Trio MRI scanner. Structural images were collected via a 3D MPRAGE T1-weighted sequence with an isotropic voxel size of 1×1×1 mm^3^, and TE/TR/TI = 2.272/2300/915 ms, FA=8. Diffusion-weighted imaging (DWI) was collected with EPI sequence with a single b-value of 1000, slice thickness of 4 mm, TE/TR = 90/5400 ms, in-plane resolution of 2×2 mm^2^, and 30 diffusion directions.

### Regions of Interest

We drew regions of interest (ROI) manually on each patient’s T1-weighted structural scan in FMRIB Software Library (FSL), using neuroanatomical landmarks because anatomical distortion made automated segmentation impractical. Representative PFC ROIs are depicted in **Figure 1.** The first slice used in PFC ROIs was one slice anterior to the corpus callosum. The anterior commissure-posterior commissure (AC-PC) line was drawn to delineate the OFC from the rest of the PFC. The OFC was drawn below the AC-PC line, anterior to the corpus callosum. The first slice used for the ACC ROI was also just anterior to the corpus callosum and included all slices where the ACC was visible. The mPFC began just anterior to the ACC and had all subsequent anterior slices. Lastly, the dlPFC included all remaining PFC; its boundaries were the OFC and ACC/mPFC. Whole thalamus ROIs were drawn based on visual differentiation of gray and white matter. MD thalamus ROIs were drawn based on the Morel Stereotactic Atlas of the Human Thalamus (Niemann et al. 2000).

**Figure 1.**
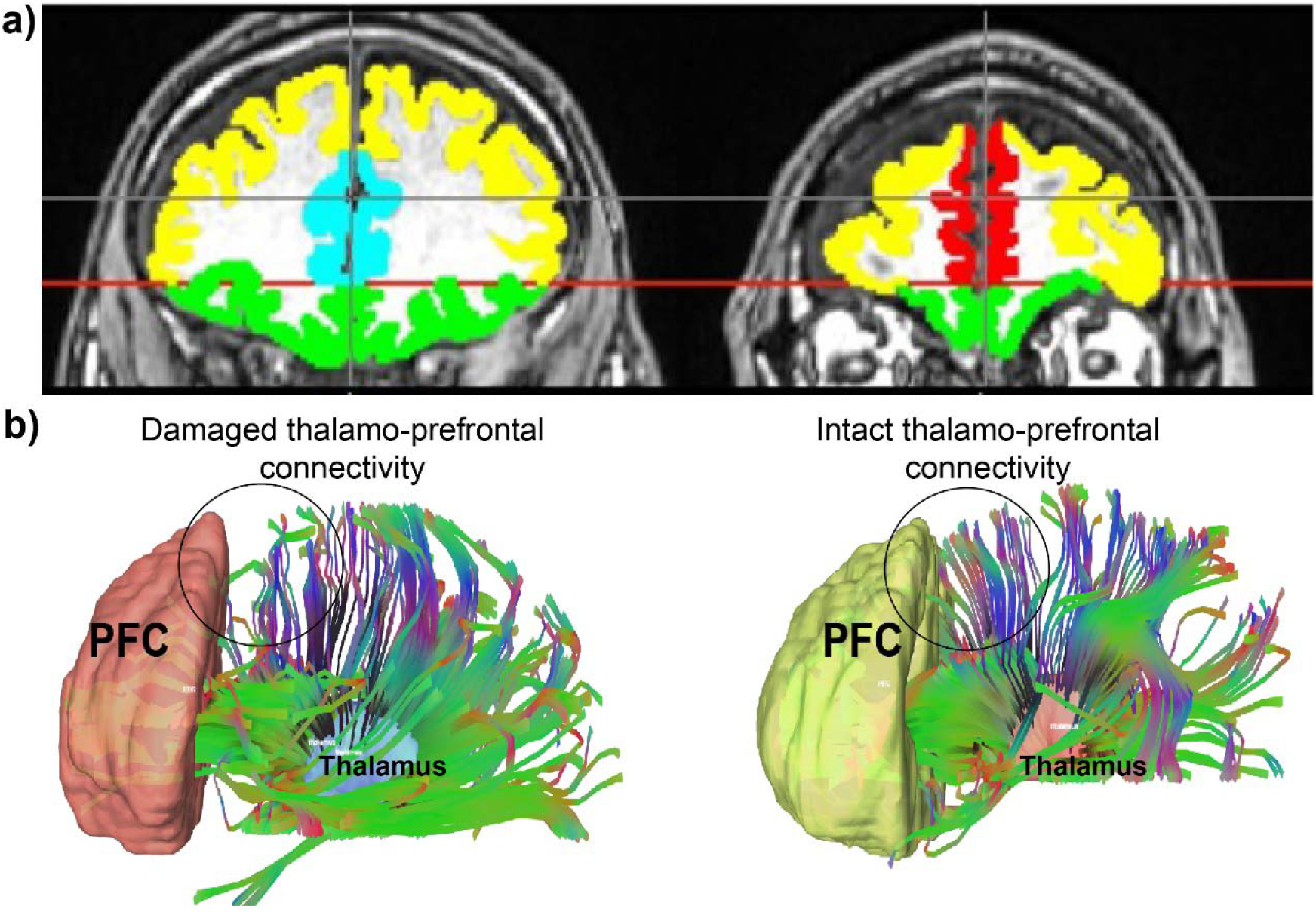
**a)** An example of PFC regions of interest with colors representing each one. Yellow = dlPFC; green = OFC; teal = ACC; red = mPFC. **b)** Thalamo-prefrontal tractography from two representative patients: one with damage to thalamo-prefrontal projections and one without. An area of decreased thalamo-prefrontal projections in the patient with damage to these connections is represented by a circle. This same area in the patient with intact projections is circled as well.

### Tractography

Diffusion MRI tractography was performed in order to characterize each patient’s thalamo-prefrontal projections. Diffusion-weighted images were corrected for the off-resonance field produced by the susceptibility distribution of the subject’s head using the FSL “topup” tool (Andersson et al. 2018), and for the eddy currents produced by the rapid switching of the diffusion gradients using the FSL “eddy” tool (Andersson and Sotiropoulos 2016). ROIs were merged to each patient’s DWI using FSL “flirt” tool and confirmed for accuracy via visual inspection. A diffusion tensor model was fitted at each voxel in the brain-extracted DWI. The DTI model was used to generate FA maps for each subject. Values for FA were calculated and compared to patient clinical outcomes. Tractography was performed using the Quantitative Anisotropy (QA) algorithm (Yeh et al. 2013), which augments deterministic tractography to correct for noisy fiber orientation distributions by incorporating anisotropic spinning along the fiber orientation. In separate experiments, the whole thalamus and MD thalamus in each hemisphere were used as seeds, and each ipsilateral prefrontal ROI was used as a target. Tractography was performed using the following tracking parameters: termination index = FA; threshold = random; angular threshold = 60°; maximum tracts = 10,000.

### Statistics

We analyzed each PFC region separately. We used a Student’s t-test to compare the mean FA values between the *Early* and *Late/Never* groups. FA values were further adjusted for age and sex using a General Linear Model (GLM).

We used linear regressions to assess clinical outcome predictability using FA values, age, and sex. Stepwise regression was performed in R using the stepAIC function. The accuracy of the models was evaluated using a leave-one-out approach.

## Results

We retrospectively analyzed patients admitted with severe TBI who underwent DWI (n = 25, 20 males, mean age 50 ± 20.9; **Table 1**). This cohort is representative of the demographics of TBI on Long Island. Three patients never regained the ability to follow commands and eventually expired, though they survived for more than 30 days. The rest of the cohort regained the ability to follow commands, ranging from two days post-injury to 178 days post-injury, with a mean and standard deviation (mean ± SD) of 34 ± 50 and a median of 12.5 days. Sixteen (64%) patients had an early return of command-following (within 30 days of injury), and nine (36%) had late or no return of command-following.

### Thalamo-prefrontal connectivity is associated with early return of command-following

We measured fractional anisotropy (FA) of white matter tracts between thalamus and dlPFC, mPFC, ACC, and OFC end regions (**Figure 1**). We performed right- and left-sided analyses separately for a total of eight separate tracts of interest. We split patients into two groups: (1) *Early* command-following group (command-following within 30 days of injury), and (2) *Late/Never* command-following group.

We found that fractional anisotropy (FA), a measure of tract integrity, was significantly lower in the *Late/Never* group for whole thalamus connectivity to left and right ACC, right mPFC, and left OFC. The mean ± SD of significantly different subregions for *Early* and *Late/Never* groups, respectively, were as follows: left ACC, 0.356 ± 0.047 and 0.307 ± 0.051; right ACC, 0.373 ± 0.05 and 0.288 ± 0.042; right mPFC, 0.367 ± 0.031 and 0.302 ± 0.034; left OFC, 0.347 ± 0.043 and 0.298 ± 0.055. The results shown are adjusted for sex and age (**Figure 2**, **Table 2**).

**Figure 2.**
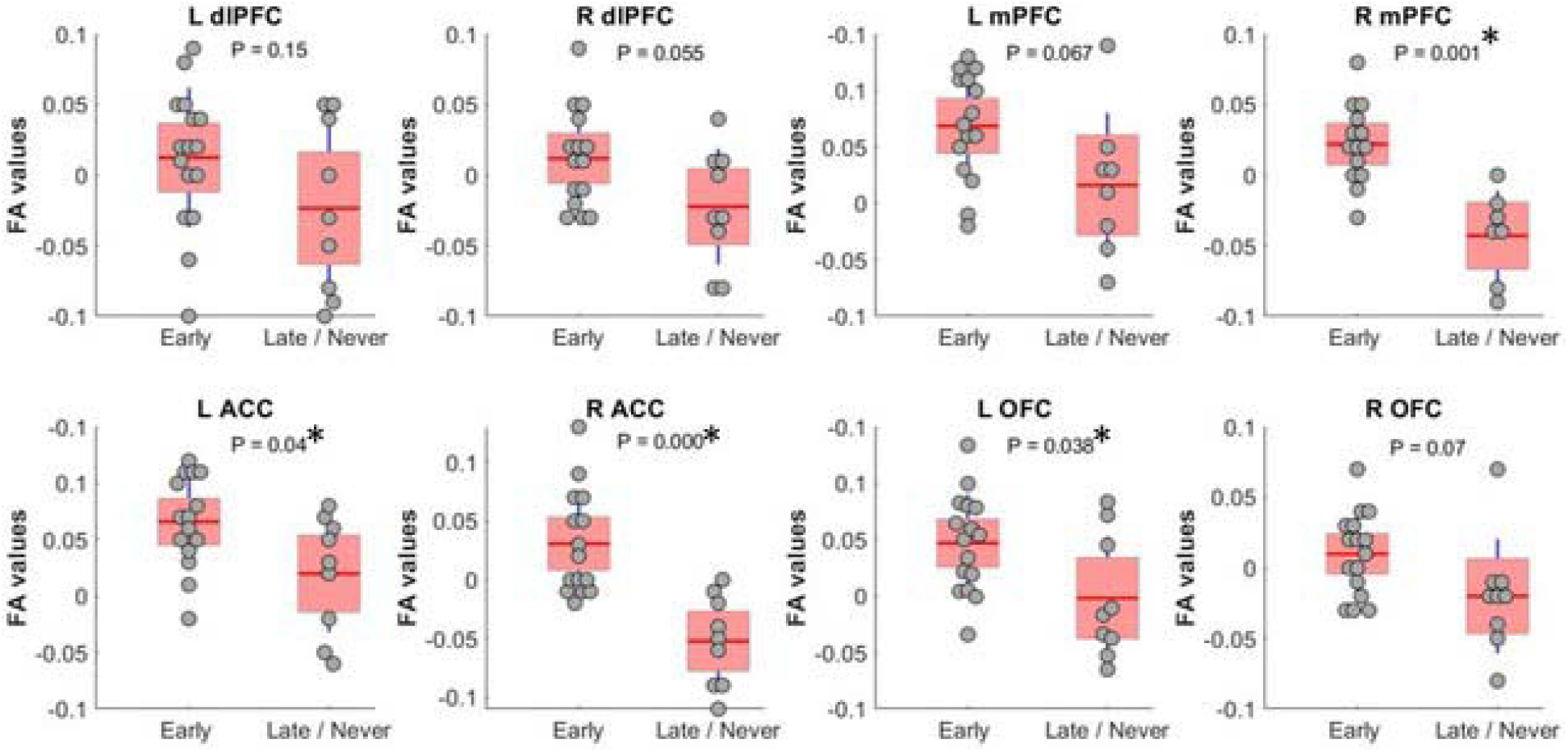
FA values (showing residual after adjusting for age and sex) between whole thalamus and PFC subregions. Subregions with significantly different FA between *Early* and *Late/Never* command-following groups are shown with an asterisk.

**Table 2.**
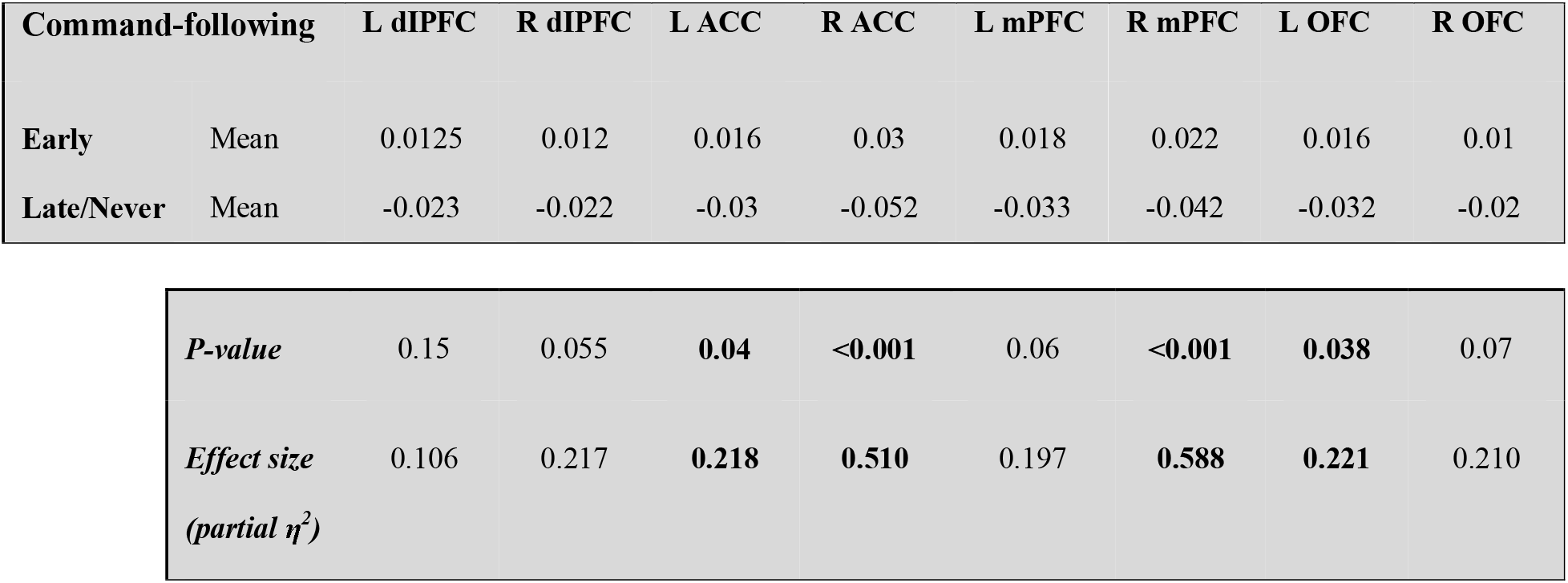
Mean, *P-values*, and effect size of FA measurements (adjusted for age and sex) between whole thalamus and PFC regions in *Early* and *Late/Never* command-following groups.

### MD-prefrontal connectivity distinguishes *Early* and *Late/Never* command-following groups

To shed light on MD’s role in command-following, as MD has the largest reciprocal connectivity to PFC, we repeated the tractography analysis using MD as the seed instead of the whole thalamus. When we restricted the analysis to MD, FA values were significantly different between *Early* and *Late/Never* groups for connectivity to right mPFC, left and right ACC, and left and right OFC. The mean ± SD FA values for each significant sample in the *Early* and *Late/Never* groups, respectively, were as follows: right mPFC, 0.343 ± 0.048 and 0.267 ± 0.031; left ACC, 0.344 ± 0.047 and 0.284 ± 0.047; right ACC, 0.334 ± 0.051 and 0.273 ± 0.046; left OFC, 0.336 ± 0.050 and 0.266 ± 0.039; right OFC 0.337 ± 0.043 and 0.282 ± 0.035. **Figure 3** shows that this result remains robust following age and sex correction using GLM. The residuals and the *P*-values for each sample after GLM analysis are presented in **Table 3**.

**Figure 3.**
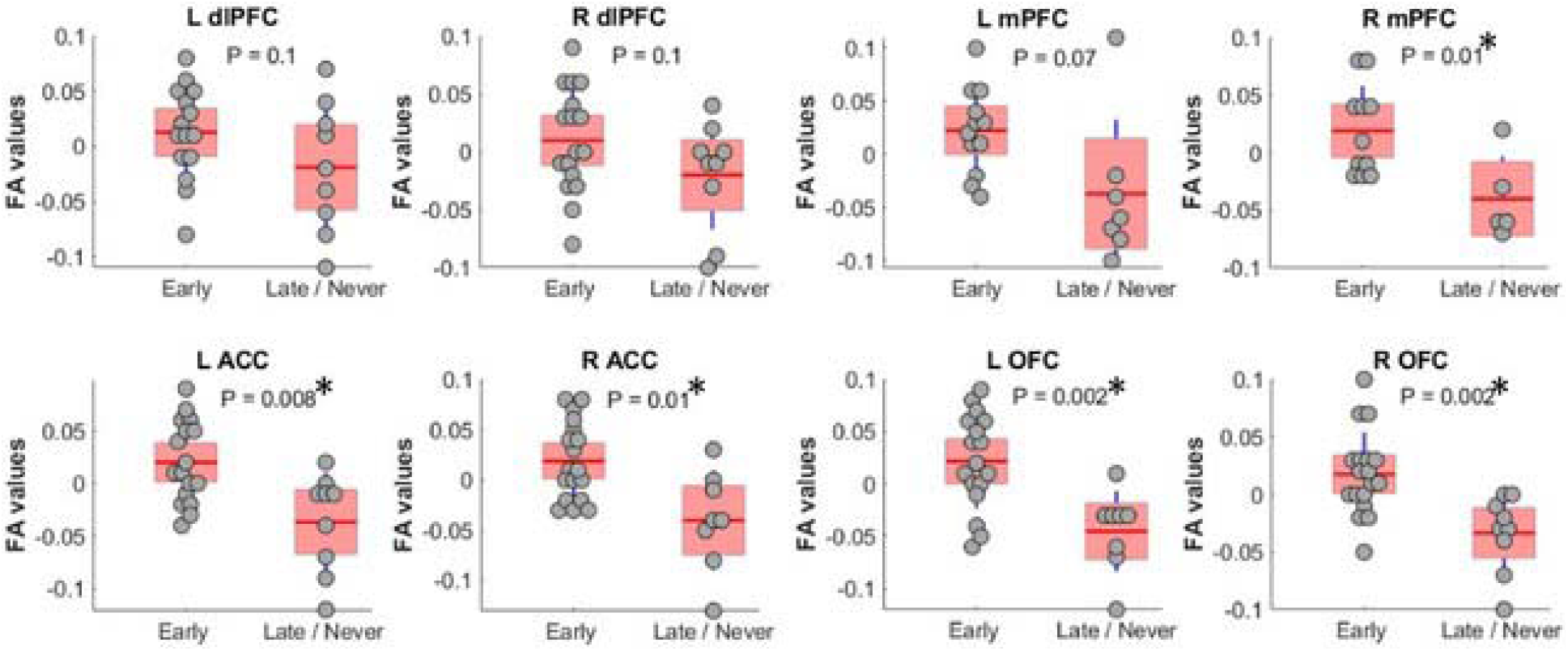
FA values (showing residual after adjusting for age and sex) between MD thalamus and PFC subregions. Subregions with significantly different FA between Early and Late/Never command-following groups are shown with an asterisk.

**Table 3.**
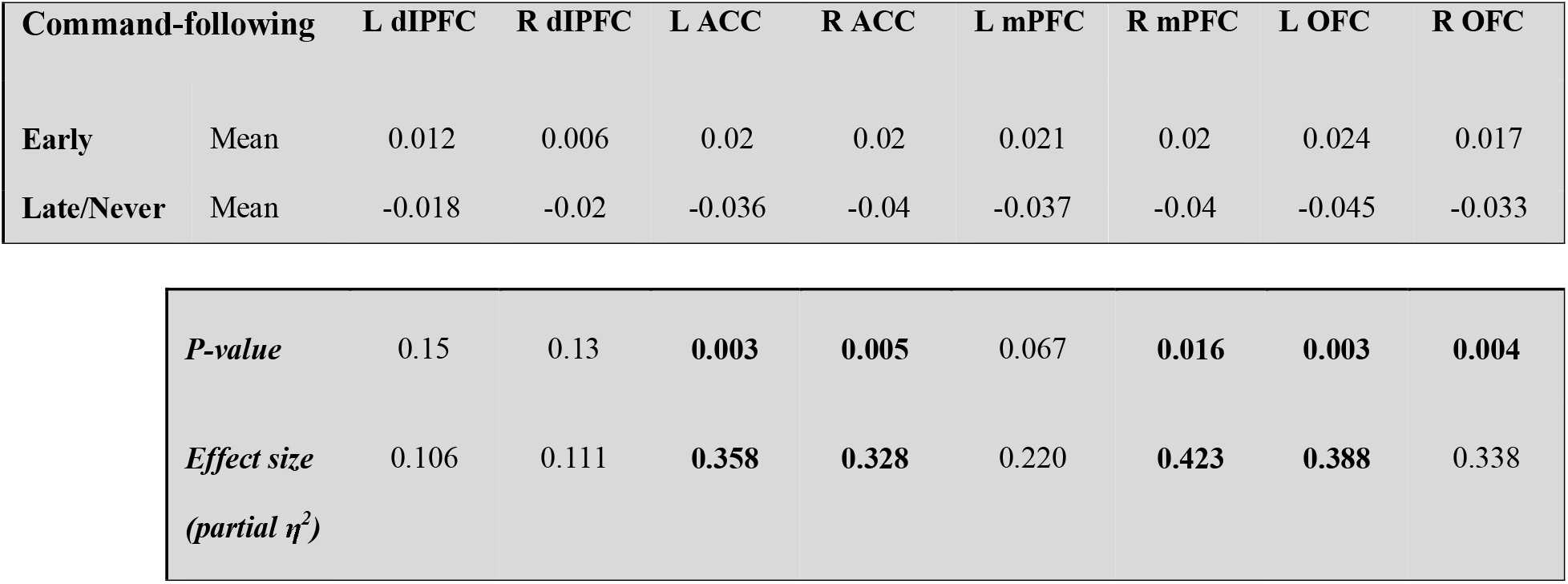
Mean, *P*-values, and effect size of FA measurements (adjusted for age and sex) between MD thalamus and PFC regions in *Early* and *Late/Never* command-following groups.

### The integrity of the thalamo-prefrontal connections predicts the return of command-following ability

We used linear regression to develop a model to predict command-following within 30 days after TBI (**Figure 4**). The model revealed that FA values for left and right ACC, mPFC, and OFC regions, along with patient sex, were predictive of patient clinical outcomes. Areas with the highest correlation in this model included left and right ACC, right mPFC, and left and right OFC. This model was further evaluated using a leave-one-out approach. The model had 72% (18/25) accuracy for predicting the ability to follow commands within 30 days of injury, with a sensitivity of 81% and specificity of 56%.

**Figure 4.**
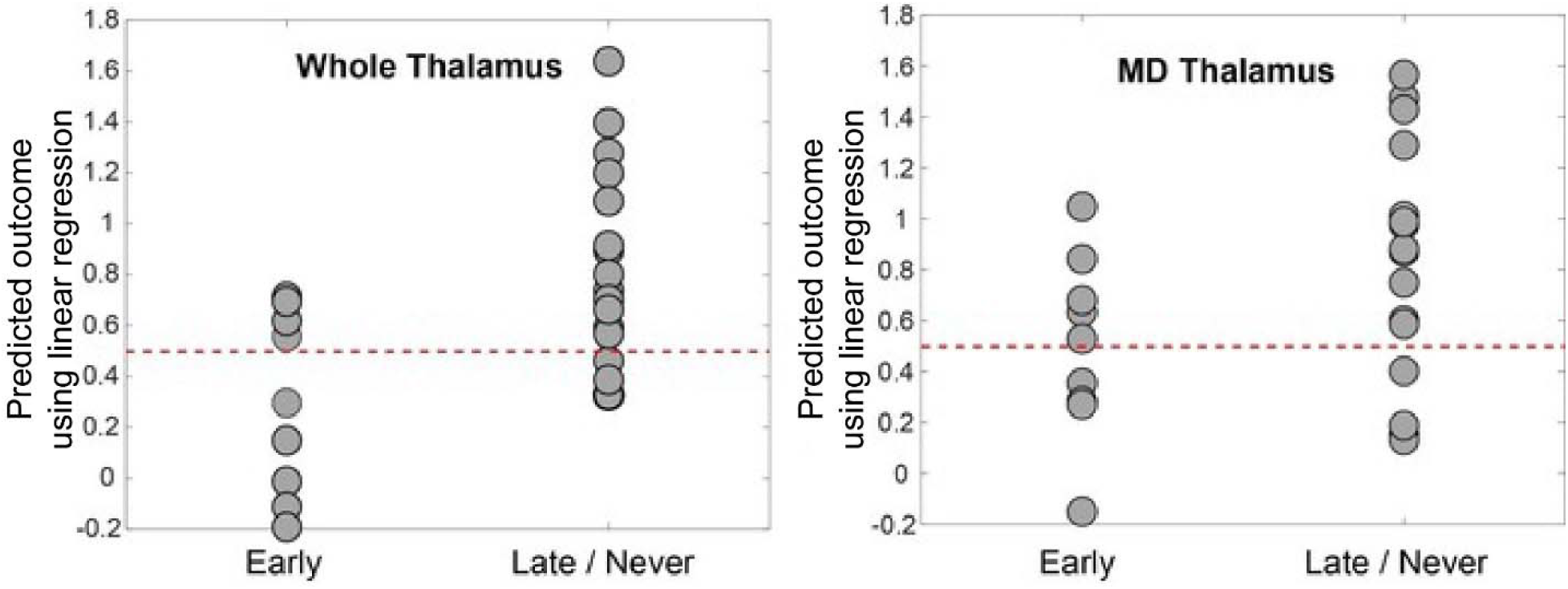
Predicted outcome by linear regression model versus actual outcome. This was generated by leave-one-out cross-validation using FA values between prefrontal regions and the whole thalamus (left panel) and MD (right panel). The dotted line at 0.5 represents the intuitive threshold used in this work to separate two different outcomes (represented by 0 and 1 in the regression). The prediction accuracy is 72% for the whole thalamus and 64% for MD. The model for whole thalamus selected left and right ACC, mPFC, and OFC regions, as well as male sex as predictive variables. The model for MD selected right dlPFC, left and right mPFC, right OFC, and male sex as predictive variables.

## Discussion

In this study, we investigated the role of the thalamo-prefrontal circuit in recovery of command-following, and thus goal-directed behavior, after TBI. We found that the integrity of thalamic connections with PFC subregions mPFC, ACC and OFC was significantly associated with early command-following. These data highlight a role for thalamo-prefrontal circuits in the recovery of goal-directed behavior after injury. Restricting our analysis to connections between MD and PFC subregions revealed similar results to the whole thalamus analysis, supporting previous research about its role in cognitively-based tasks (Alexander and Fuster 1973; Dolleman-van der Weel, Morris, and Witter 2009; Peräkylä et al. 2017; Schmitt et al. 2017; Sébastien Parnaudeau, Bolkan, and Kellendonk 2018; Rikhye, Gilra, and Halassa 2018). Our data stand in contrast to recently published studies in which thalamic injury was not correlated with unconsciousness (Hindman et al. 2018; Rohaut et al. 2019). However, these studies did not consider thalamocortical connectivity nor use the detailed anatomical analysis of our research. Moreover, although recently proposed models de-emphasize the frontal lobes’ role in consciousness (Koch et al. 2016), our data show that PFC, more specifically its medial subregions, is important for recovery after TBI.

An important finding of our study is the apparently critical role of medial portions of PFC (including mPFC and ACC). The mPFC is part of the default mode network (DMN), which governs the resting state of consciousness (Raichle et al. 2001; Greicius et al. 2003). Previous studies have demonstrated that DMN connectivity is decreased at lower levels of consciousness such as the vegetative and minimally conscious states (Vanhaudenhuyse et al. 2010; Fernández-Espejo et al. 2012). Another possible explanation is that medial PFC activity is correlated to task performance during periods when dlPFC activity is weak (Silton et al. 2010), as in the low-arousal state that characterizes the period after TBI. Moreover, task-related rules engage specific patterns of activity in PFC. Recent animal studies have shown that cortical representations needed for task performance require thalamic support (from higher-order thalamic nuclei such as MD) to be sustained (Schmitt et al. 2017). Thus, injuries that extend to the thalamus and thalamocortical projections likely hinder the formation and maintenance of these cortical representations associated with different goal-directed behaviors and delay command-following in TBI patients. In agreement with prior research (Schiff et al. 2007; Cain et al. 2021), our data suggest that augmenting activity in the thalamus, particularly MD, through techniques such as neurostimulation may improve goal-directed behavior after TBI.

### Study Limitations

Some limitations do affect our results. The study cohort was a convenience sample of severe TBI patients, which contributed some variability in demographics, injury patterns, and clinical management. However, given the heterogeneity of this population, it is not possible to make progress in TBI research without investigations of cohorts that reflect “real-world” TBI demographics. Furthermore, the MRI imaging data in this study were acquired for clinical care, and were not fully optimized for research; nonetheless, we consider the imaging findings robust to multiple comparisons corrections. A prospective imaging study is warranted to confirm the findings in this study. Together, our data robustly highlight the role of thalamo-prefrontal connectivity in command-following.

## Author contributions

Study concept and design: SM and CBM. Study supervision and coordination: CBM, CH, and SM. Drafting/revising the manuscript for content, including medical writing for content: MEC, JRS, CBM, SS, CH, SM. Acquisition of data: PLS and MEC. Analysis or interpretation of data: MEC, PLS, LA, ZW, CBM, CH, SM. Statistical analysis: SM, CH, PS. Obtaining funding: CBM and SM.

## Acknowledgments

The authors would like to thank Anthony Asencio, Susan Fiore, and Dr. Raphael Davis for their generous contributions to the conduct of the study. We also thank the Department of Neurosurgery at Stony Brook University Hospital for supporting this research.

## Data Availability

The data for this study are available upon reasonable request.

